# A biophysical rationale for the selective inhibition of PTP1B over TCPTP by nonpolar terpenoids

**DOI:** 10.1101/2023.04.17.537234

**Authors:** Anika J. Friedman, Hannah M. Padgette, Levi Kramer, Evan T. Liechty, Gregory W. Donovan, Jerome M. Fox, Michael R. Shirts

## Abstract

Protein tyrosine phosphatases (PTPs) are emerging drug targets for many diseases, including type 2 diabetes, obesity, and cancer. However, a high degree of structural similarity between the catalytic domains of these enzymes has made the development of selective pharmacological inhibitors an enormous challenge. Our previous research uncovered two unfunctionalized terpenoid inhibitors that selectively inhibit PTP1B over TCPTP, two PTPs with high sequence conservation. Here, we use molecular modeling with experimental validation to study the molecular basis of this unusual selectivity. Molecular dynamics (MD) simulations indicate that PTP1B and TCPTP contain a conserved h-bond network that connects the active site to a distal allosteric pocket; this network stabilizes the closed conformation of the catalytically influential WPD loop, which it links to the L–11 loop and *α*3 and *α*7 helices—the C-terminal side of the catalytic domain. Terpenoid binding to either of two proximal allosteric sites—an *α* site and a *β* site—can disrupt the allosteric network. Interestingly, binding to the *α* site forms a stable complex with only PTP1B; in TCPTP, where two charged residues disfavor binding at the *α* site, the terpenoids bind to the *β* site, which is conserved between the two proteins. Our findings indicate that minor amino acid differences at the poorly conserved *α* site enable selective binding, a property that might be enhanced with chemical elaboration, and illustrate, more broadly, how minor differences in the conservation of neighboring—yet functionally similar—allosteric sites can have very different implications for inhibitor selectivity.

## Introduction

An important challenge in the development of pharmaceutical inhibitors is the joint optimization of both binding affinity and selectivity. For example, a medicinal chemist might increase the binding affinity of a compound for a drug target by increasing its nonpolar surface area, but large nonpolar compounds often bind nonspecifically, a source of toxic side effects (*1, 2*). Our previous work suggests that some nonpolar terpenoids can exhibit selective interactions with protein tyrosine phosphatases (PTPs) (*3, 4*). Of course, nonpolar terpenoids are not generally useful as drugs; they have questionable metabolic stability and low solubility, which limits bioavailability (*5*). Nonetheless, they are promising scaffolds for drug development. Paclitaxel and artemisinin are notable examples of highly efficacious terpenoid-based therapeutics (*6, 7*). This study uses selective terpenoid inhibitors of PTPs to study the molecular basis of selective interactions between nonpolar compounds and proteins—a variety of interactions that remains challenging to exploit in rational drug design.

The human genome contains 107 PTPs, of which 38 are tyrosine-specific—or “classical”—PTPs that share a highly conserved catalytic domain (*8*). Pronounced structural conservation at the active site of these PTPs has made the development of selective inhibitors difficult and driven researchers to focus on less conserved allosteric sites (*9, 10*). PTP1B provides a prominent example. This enzyme is a longstanding target for treating type 2 diabetes, obesity, and HER2+ breast cancer (*11 –18*) and has emerged more recently as a promising immuno-oncology target (*19*). Its structural similarity to T-cell PTP (TCPTP), which has 80% sequence homology, has focused medicinal chemistry efforts on a C-terminal allosteric site, which can enhance selectivity. Intriguingly, even this site has relatively high structural conservation, which enhances the difficulty of developing selective compounds. A detailed understanding of selective allosteric inhibition mechanisms could yield new approaches to improve selectivity.

The promise of PTP1B as a drug target has motivated the development of an extraordinarily broad set of inhibitors. Most of these compounds bind to the active site, but several bind to the allosteric site. The most well-studied allosteric inhibitor is the benzofuran 3-(3,5-Dibromo-4-hydroxy-benzoyl)-2-ethyl-benzofuran-6-sulfonicacid-(4-(thiazol-2-ylsulfamyl)-phenyl)-amide, hereafter referred to as BBR, which was discovered in an early highthroughput screen and has comparable selectivity to other potent PTP1B inhibitors (*20, 21*). Upon binding to PTP1B, BBR is thought to engage in both *π*-stacking interactions with a non-conserved PHE280 residue (*22*) and h-bonds with neighboring residues in the C-terminal allosteric pocket. Using x-ray crystallography, MD simulations, and mutational analysis, we previously demonstrated that BBR and the small terpenoid hydrocarbon amorphadiene (AD) disrupt the allosteric network in PTP1B in a similar manner(*3*). AD and the chemically similar compound *α*-bisabolene (AB) are surprisingly selective for PTP1B, which they inhibit 5–7X more potently than TCPTP, a selectivity similar to BBR, which inhibits PTP1B 5–6X more potently (*4, 20*). In general, AD and AB are intriguing inhibitors because they cannot form h-bonds or other specific interactions and must therefore exhibit a mechanism of selectivity distinct from that of BBR. Understanding the mechanism by which non-functionalized inhibitors achieve both moderate potency and isoform selectivity could inform the optimization of allosteric inhibitors for both PTP1B and TCPTP.

In prior work, we used molecular modeling to study how AD inhibits PTP1B. Our simulations showed that AD can sample two neighboring regions of the C-terminal allosteric site, both of which require a disordered *α*7 helix to form a stable complex (*3*). In forming this complex, AD disrupts a h-bonding network that stabilizes closure of the catalytically essential WPD loop. This previous modeling work helped explain the mechanisms of AD binding and inhibition, but it did not shed light on the molecular basis of its selectivity for PTP1B over TCPTP, which has sequence differences in both the *α*6 and *α*7 helices, but not in the *α*3 helix, the residues in the putative h-bond network, or active site.

In the present study, we used molecular modeling to study the mechanistic basis of inhibitor selectivity for AD and AB, along with some experimental testing of these predictions. This work seeks to understand how minor differences in the sequences of PTP1B and TCPTP might cause differences in either binding affinity or allosteric modulation. The inclusion of a second terpenoid inhibitor helps us test—and expand on—our previously proposed mechanism of selectivity. The direct comparison of binding to two PTPs helps us answer several pressing questions:

- Does an analogous h-bonding network stabilize the closed conformation of the WPD loop in both PTP1B and TCPTP? Can the network in TCPTP be modulated through the disruption of the *α*3–*α*7 interface in a PTP1B-equivalent manner or is a different interface disrupted?
- Do non-functionalized terpenoids such as AD and AB exhibit a conserved mechanism of binding to and inhibition of PTP1B? Does this mechanism also apply to TCPTP?
- Are analogous *π*-stacking interactions with PHE280 or other residues present with inhibitor binding to TCPTP as they are with PTP1B?

### Materials and Methods

#### Molecular Dynamics (MD) Simulations

We prepared PTP1B and TCPTP for MD simulations by starting with four X-ray crystal structures: apo PTP1B (PDB code: 1SUG), apo TCPTP with the WPD loop closed and an ordered *α*7 helix (PDB code 7F5N), apo TCPTP with the the WPD loop open and a disordered-unresolved *α*7 helix(PDB code 1L8K), and PTP1B in complex with AD (PDB code: 6W30) were taken from the Protein Data Bank for use as initial conformations after post-processing (*4, 23 –25*). For each structure, we removed crystallized waters, glycerol, and Mg^2+^, adjusted the protonation state to a pH of 7 using the H++ web-server, added Na^+^ ions to neutralize the net charge, and hydrated the protein with a TIP3P water box, maintaining a minimum distance of 10 Å between the protein or ligand and the periodic boundary. Given routine incorrect predictions by H++, the catalytic CYS215 residues were manually verified to be in the expected deprotonated state for physiological pH conditions.

We carried out MD simulations with GROMACS 2020.4 (*26*) on the Bridges-2 cluster at the Pittsburgh Supercomputing Center. In all simulations, we modeled PTP1B with the AMBER *ff99sb-ildn* force field and parameterized AD and AB with the Open Force Field v.1.3.0 “Parsley” (*27*). All analysis scripts and input parameters can be found in the repository at https://github.com/shirtsgroup/TCPTP. Ligand parameterization scripts can be found in repository folder “Ligand Parameters”. We carried out an energy minimization to 100 kJ/mol/nm force tolerance, and equilibrated the protein in the NVT ensemble at 300 K for 100 ps, followed by equilibration to the NPT ensemble at 300 K and 1 atm for 100 ps. All simulations used the velocity rescaling thermostat (*28*) and Beredensen weak-coupling barostat. Further configuration details for the simulations appear in the repository folder “data/mdp”. We ran all MD simulations for 300 ns (unrestrained NPT) and visualized using PyMOL 2.4 (*29*).

X-ray crystal structures of PTP1B and TCPTP with the WPD loop in an open conformation both apo or ligand bound have a disordered *α*7 helix which prevents resolution in the crystal structure. Generating disordered *α*7 helix conformations for PTP1B was done according to the procedure explained in our previous study (*3*). The same general method was utilized to generate the disordered TCPTP conformations; however, the *α*7 helix on TCPTP was more resistant to destabilization. For the restrained heating method of generating disordered conformations described in that paper, the temperature was still increased from 400 to 500 K over 300 ns, but an additional 100 ns at 500 K was necessary to disorder the *α*7 helix to below 50% *α* helicity. *α* helicity is defined as the percent of residues in the *α*7 helix (residues 284–294 for TCPTP) which are in the *α* helical conformation as defined by the Defined Secondary Structure Prediction (DSSP) algorithm. Three disordered conformations were selected from the final 50 ns of this trajectory. For the BBR-bound method, when BBR was in complex with PTP1B the *α*7 helix disordered within 50 ns, but when bound to TCPTP simulations were run for 300 ns and the *α*7 helix stabilized at *∼*50% *α* helicity. Two partially disordered conformations were selected from the final 50 ns of the 300 ns trajectory.

There is no available crystal structure for TCPTP in complex with either AD or AB binding poses; for both the initial configurations were determined using molecular docking and through PTP1B-AD structure alignment. AutoDock Vina 1.2.0 (*30*) was used to perform molecular docking to apo TCPTP structures with a disordered *α*7 helix and open WPD loop. For AD docking, the TCPTP structure used was the centroid from clustering on backbone atoms of the apo trajectory of all five generated disordered *α*7 helix configurations.

Docking was performed for each of the five structures with a 27 nm^3^ search space centered on the crystal binding site of AD to PTP1B, as we were originally operating under the assumption that the ligands would bind to approximately the same location on TCPTP. For docking, a search exhaustiveness of 32 was utilized and the 15 highest affinity binding modes were generated. Of these 15 modes, all those determined to be distinct and unlikely to be sampled in the same trajectory were chosen as initial configurations. Distinct conformations were determined based on their proximity to one another as those whose ligand center of mass was within 5 Å are likely to be sampled within the same trajectory given the COM RMSD of AD is approximately 7 Å. This procedure produced 4–5 binding locations, depending on the TCPTP structure used. For AB, only the two disordered *α*7 helix conformations which lead to stable AD binding in at least one initial configuration were used for docking otherwise the procedure was identical.

None of the initial binding poses generated by docking were in the crystal binding location for PTP1B and thus an additional configuration was added with each ligand placed in the AD crystal binding location as determined by aligning the protein backbone. Minor modifications were made (movements of <1 Å) to account for steric clashes between the ligand and protein which produced infinite energies during minimization. All initial configurations were initially run for 50 ns unrestrained NPT following equilibration in order to determine if the ligand was stable. Any binding pose in which the ligand fully dissociated from the protein was not continued for the full 300 ns.

A similar procedure was used to generate starting binding configurations for PTP1B-AB complex since there is no available crystal structure for AB in complex with PTP1B due to low solubility. For AB docking, the PTP1B structure used had a WPD loop in the open conformation and a disordered *α*7 helix. Both the originally generated structure and the centroid from clustering on backbone atoms of an apo trajectory were used to generate docking configurations. For each of the two structures, docking was performed in a 9 nm^3^ search space centered on the crystal binding site of AD to PTP1B, since previous experimental results indicated a conserved binding location (*4*). These search space dimensions adequately encompassed the allosteric site of PTP1B, as the maximum COM RMSD achieved by AD in complex with PTP1B was *>* 1 nm and this volume encompassed the entirety of the surfaces presented by the *α*3, *α*6, and *α*7 helices. The 20 highest affinity binding modes were generated for each PTP1B configuration used and, as above, all those determined to be distinct were chosen as initial configurations for MD simulations. This resulted in 9–11 binding locations, depending on the PTP1B structure used for docking. As above, all binding poses in which the ligand fully dissociated from PTP1B during the initial 50 ns simulations were not continued for a full 300 ns trajectory.

### Analysis of MD Trajectories

Before completing analysis on our MD trajectories in detail, we carried out two important processing steps: (i) removal of correlated trajectory frames (ii) removal of unequilibrated trajectory frames and determination of convergence. Correlated trajectory frames were removed with ruptures 1.1.6 (*31*) and unequilibrated trajectory frames were removed based on the root-mean-square deviation (RMSD) of backbone atoms, relative to the starting structure for the production simulation (further details in previous work (*3*)). Following these processing steps the number of uncorrelated frames per nanosecond range from 2–4 depending on the trajectory. This process could prevent the resolution of rare events; however, all data measurements were verified to not have a strong effect on the mean values obtained and simply increase the accuracy of statistical measurements.

AD and AB exhibited several distinct binding conformations when bound to PTP1B compared to binding with TCPTP. In complex with PTP1B, both ligands bound to what we refer to as the *α* site (Figure 2A). The *α* site is defined as simultaneous contacts with the *α*3 and *α*7 helices. The *α* site encompasses both the loc1 and loc2 identified in our previous study as ligand oscillation frequency between the two sites was so high the two sites are unlikely to be kinetically distinct (*3*). In contrast, in complex with TCPTP, both ligands bind to the *β* site, which is defined as simultaneous contacts with three structures: at least of the two *β* sheets in *β*8–10, as well as the *α*3 helix (Figure 2B). The *β* site was identified from the trajectories of TCPTP in complex with both AD and AB. Of the 23 initial configuration simulated for the TCPTP-AD complex only 6 remained in contact with the protein after the 50 ns evaluation simulation. Of these 6, two were bound in the region classified as the *β* site and only one of the two was initiated near this site, one was bound to the *α* site before becoming unstable after approximately 150 ns, and the remaining three sampled multiple binding locations but ended the trajectory dissociated from the protein. Of the 8 initial configurations simulated for the TCPTP-AB complex 5 remained bound to the protein after the 50 ns evaluation simulation. Of these 5, three were bound in the region classified as the *β* site (only one was initiated there) and the other two sampled distinct alternate binding positions near the *α*4 helix. From the complex of TCPTP with AD and AB the only conserved and repeatable binding location was the identified *β* site. This site is also identical to loc4 identified in our previous study from binding of AD to PTP1B with a truncated *α*7 helix and it is overlapping with the 197-site identified by Keedy et al. (*32*) using a high throughput fragment library search.

Both PTP1B and TCPTP have catalytically active and inactive states, in which the active conformation has a closed WPD loop and ordered *α*7 helix while the inactive conformation has an open WPD loop and disordered *α*7 helix. We classified the WPD loop conformation by the distance between the *α* carbons of D181 and C215 for PTP1B and D182 and C216 for TCPTP (i.e., the catalytic acid and the nucleophile, respectively) (*33 –35*), as measured using the compute_distances function of MDTraj. A distance of >1 Å was defined as open and otherwise the loop was defined as closed. This metric was explored and validated in our previous work (*3*). The helicity of the *α*7 helix in our MD trajectories was quantified using the DSSP algorithm implemented in MDTraj 1.9.4 (*36*). This algorithm characterizes the secondary structure of each residue based on the *ϕ* and *ψ* torsional angles. This analysis allowed us to characterize the order, or lack thereof, of the *α*7 helix.

To further classify the structure of PTP1B throughout the simulations, we evaluated the RMSD of the backbone atoms and the root-mean-square-fluctuation (RMSF) of select protein regions, relative to a centroid structure. We defined the centroid structure by clustering each trajectory on the backbone atoms of the equilibrated trajectory using gmx_cluster and taking the centroid of the most populated cluster (consistently containing >90% of the trajectory). For ligand bound trajectories, the center-of-mass (COM) RMSD for the ligand was also computed using bootstrapping on the uncorrelated configurations to determine the mean and standard error of the ligand COM RMSD value.

The catalytic domain of PTP1B has seven *α* helices, several of which play important roles in allosteric communication. We quantified inter-helical interactions and helix-ligand interactions between these influential helices as those with a residue-residue or residue-ligand distance of less than 4 Å. We defined inter-helical interactions disrupted by ligand binding as those that occur significantly less (*p <* 0.05) in the ligand bound vs. corresponding apo conformation. We calculated the p-value using Welch’s T-test for the fraction of the simulation time that the interaction was present for the ligand bound (AD or AB) compared to apo trajectories. Additionally, *π*-stacking interactions between the protein and ligand were defined as interactions in which at least two adjacent sp^2^ carbons in the protein are a distance of less than 3.5 Å from two adjacent sp^2^ carbons on the ligand.

We isolated allosterically influential h-bonds with several steps. (i) We used the Baker-Hubbard model implemented within MDTraj to identify h-bonds. This model uses a proton donor-acceptor distance of 2.5 Å and a donor-acceptor angle of less than 120° to classify h-bonds. (ii) We removed h-bonds formed in a majority of all trajectories, regardless of WPD loop conformation or the presence of an allosteric inhibitor, or formed between adjacent (within 3) residues and calculated the percent of the trajectory in which each of the remaining bonds appeared. (iii) For each h-bond, we determined the mean frequency formed for both apo WPD_open_, apo WPD_closed_. (iv) We identified bonds that showed a statistically significant (*p <* 0.01) difference between the groups. (v) Using our statistical threshold, we selected bonds that appeared more in either apo WPD_open_ or apo WPD_closed_ (with a minimum appearance of 70% in their primary state) to define h-bonding networks in each of these conformations. Notably, no h-bonds appeared significantly more or less frequently (given the above selection criteria) with ligands bound than in the apo WPDopen state.

### Computational Mutation Generation

A variety of mutations were made computationally to both PTP1B and TCPTP for validation of the simulation approaches by experiment. All mutant simulations involved the same protocol. Each mutation was created from the centroid of the most populated cluster obtained from the apo protein trajectory clustered on protein backbone atoms.

The mutation itself was performed using Modeller 10.1 then a 50 ns apo simulation was completed with each mutant in order to stabilize the structure. For TCPTP mutants, AD and AB were then placed in three different binding conformations all within the defined *β* site as described above. These three conformations were centroid structures from the trajectories with AD or AB bound to TCPTP. One trajectory was chosen at random and the centroid of the trajectory was chosen as references. The other two centroids were from those trajectories which had the highest heavy-atom RMSD compared to the reference centroid to get a diversity of binding locations. For PTP1B mutants, AD and AB were placed using the same procedure but with binding sites within the previously defined *α* site. 300 ns simulations were then performed of each mutant complex and the time in nanoseconds for which the ligand remained bound to the defined site was evaluated and called the ligand retention time. The ligand COM RMSD was only calculated for the portion of the trajectory in which the ligand was bound to the *α* site or *β* site for PTP1B and TCPTP respectively.

### Experimental Methods

#### Materials

We purchased yeast extract, sodium chloride, LB broth (Miller), potassium phosphate monobasic and dibasic, tris base, tetracycline hydrochloride, magnesium sulfate heptahydrate, imidazole, 4-(2-hydroxyethyl)-1-piperazineethanesulfonic acid) (HEPES), and pre-made 1M HEPES buffer (pH 7.3) from Fisher; Triton X-100, tris(2-carboxyethyl)phosphine (TCEP), phenylmethylsulfonyl fluoride (PMSF), bovine serum albumin (BSA), and p-nitrophenyl phosphate (pNPP), dimethyl sulfoxide (DMSO) from MilliporeSigma; Phusion and DNase I from New England Biolabs; agar and M9 salts from Becton Dickinson; glucose and Nalpha-4-Tosyl-L-arginine methyl ester hydrochloride (TAME) from Acros Organics; tryptone from Research Products International; isopropyl *β* D-1-thiogalactopyranoside (IPTG) from ChemCruz; kanamycin sulfate from IBI Scientific; carbenicillin from Gemini Bioproducts; lysozyme from Alfa Aesar; 10 kDa spin columns from Sartorius; 4-20% Criterion TGX stain-free protein gels from Bio-Rad; and HisTrap and HiTrap columns from GE Healthcare.

***E. coli* Strains** We used chemically competent NEB stable (#C3040H) cells for cloning and NEB BL21(DE3) (#C2527H) for protein overexpression.

**Cloning and Molecular Biology** We constructed plasmids with Gibson assembly (50 ° C for 1 hour). Table S1 lists gene sources, and Table S2 lists primers used for site-directed mutagenesis.

### Protein expression and purification

We overexpressed all proteins examined in this study in BL21(DE3) cells by carrying out steps described previously: (i) We used Gibson assembly to introduce mutations for PTP1B or TCPTP encoded by a pET16b vector and transformed sequence-confirmed plasmids into BL21(DE3) cells. (ii) We used an individual colony from each transformation to inoculate 20 mL of LB media, which we incubated at 37°C in an incubator-shaker (225 rpm) for 6 hours, prior to inoculating 1 L of rich induction media (20 g tryptone, 10 g yeast extract, 5 g sodium chloride, 50 *μ*g/mL carbenicillin, 72 mL 5X M9 salts solution, 20 mL of 20% glucose solution). (iii) We grew the resulting inoculum at 37°C and 225 rpm. (iv) At an OD600 of 0.5–0.8, we added 500 *μ*M IPTG to induce protein expression and incubated the induced flasks at 22°C and 225 rpm for 18-20 hours. (v) We pelleted the final culture (5000 rpm for 10 minutes in a Beckman J2-HS floor centrifuge), disposed of the supernatants, and stored the cell pellets at -80°C freezer for future purification. We purified PTP variants with fast protein liquid chromatography (FPLC) using previously described protocols.

In brief, we lysed cells with chemical lysis buffer (for each gram of cell pellet, we used 4 mL of 20 mM tris base, 50 mM sodium chloride, 1% Triton X-100, and pH 7.5 supplemented with 2 mg MgSO4-7H2O, 2 mg TAME, 0.5 mL TCEP (0.5 mM), 3.75 *μ*L PMSF solution, 1 mg lysozyme, and 30 U DNase), removed protein debris with saturated ammonium sulfate (20% for PTP1B variants and 10% TCPTP variants, respectively), and extracted the supernatant. We purified PTPs by using nickel affinity chromatography (HisTrap HP with 50 mM Tris-HCl, 300 mM sodium chloride, and 0.5 mM TCEP at pH 7.5 with and without 500 mM imidazole) followed by anion exchange chromatography (HiTrap HP 5 mL with 50 mM HEPES, and 0.5 mM TCEP at pH 7.5 with and without 1 M NaCl). We concentrated final protein fractions into NaCl-free anion exchange buffer (10,000 kDa, Sartorius) and stored them in 20% glycerol at -80°C.

### Enzyme kinetics

We characterized the activity of PTP variants on p-nitrophenyl phosphate (pNPP). In brief, we used 96-well plates to prepare 200-*μ*L reactions with 50 nM enzyme, 50 *μ*g/mL BSA, and varying amorphadiene concentrations (0.5 – 500 *μ*M) in 50 mM HEPES buffer (pH = 7.3) with 10% DMSO. We used 20 mM substrate to initialize the reactions and monitored the formation of p-nitrophenol (pNP) by measuring absorbance at 405 nm at 30-s intervals over 8 minutes (SpectraMax iD3 plate reader). We used standard curves to convert absorbance measurements to product concentrations and used the slopes of the initial rate regime for Vo.

## Results

### Allosteric networks are similar between the PTPs

MD simulations indicate that PTP1B and TCPTP possess functionally very similar allosteric systems (Figure 1A). Both proteins contain a conserved network of h-bonds that connects the active site (P-loop and WPD-loop) to the allosteric site (L–11 loop, *α*3, *α*6, and *α*7 helices). These h-bonds bonds are present more frequently (p < 0.05) in the apo closed state than in the open state, an indication that they stabilize closure of the WPD loop (Figure S6). In addition to their shared h-bonds, both networks also include several distinct peripheral bonds. For example, TCPTP has two additional h-bonds between the P-loop and adjacent residues, and PTP1B has one between the *α*6 and *α*1 helices (Figure S7)(*3*). Though these bonds are present less often in the open than closed conformations of both proteins, their location and lack of connectivity to other bonds in the network suggests they do not contribute to allosteric communication between the active and allosteric sites.

**Figure 1:**
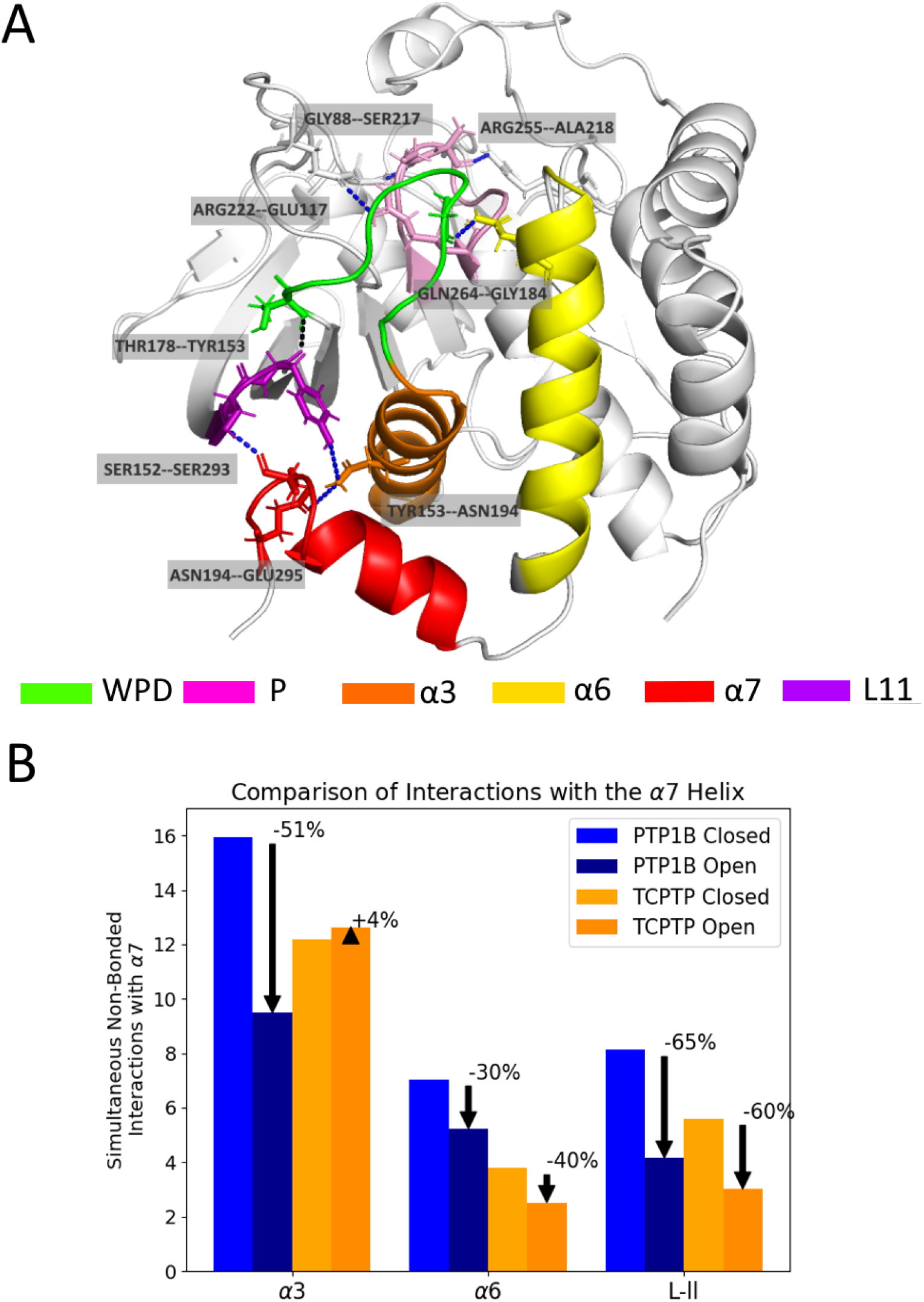
TCPTP possesses a PTP1B-like allosteric network. (A) In TCPTP, a h-bond network connects the WPD loop to the allosteric site. Each bond in this network involves residues homologous to those found in PTP1B (*3*). (B) Disordering of the *α*7 helix in PTP1B and TCPTP disrupts several additional nonbonded interactions. This figure shows the number of non-bonded interactions present between the *α*7 helix and the *α*3, *α*6, and L–11 loop when the WPD loop is open and closed (when the *α*7 helix disordered or ordered, respectively). The percentages reflect the percent difference in the number of interactions formed in the closed state over the open state. For PTP1B, the transition between these states disrupts the *α*3– *α*7 interface, followed by the L–11–*α*7 interface; for TCPTP, disruption is largely localized to non-bonded interactions at the L–11–*α*7 interface. See figure S10 for a diagram of disrupted interactions.

**Figure 2:**
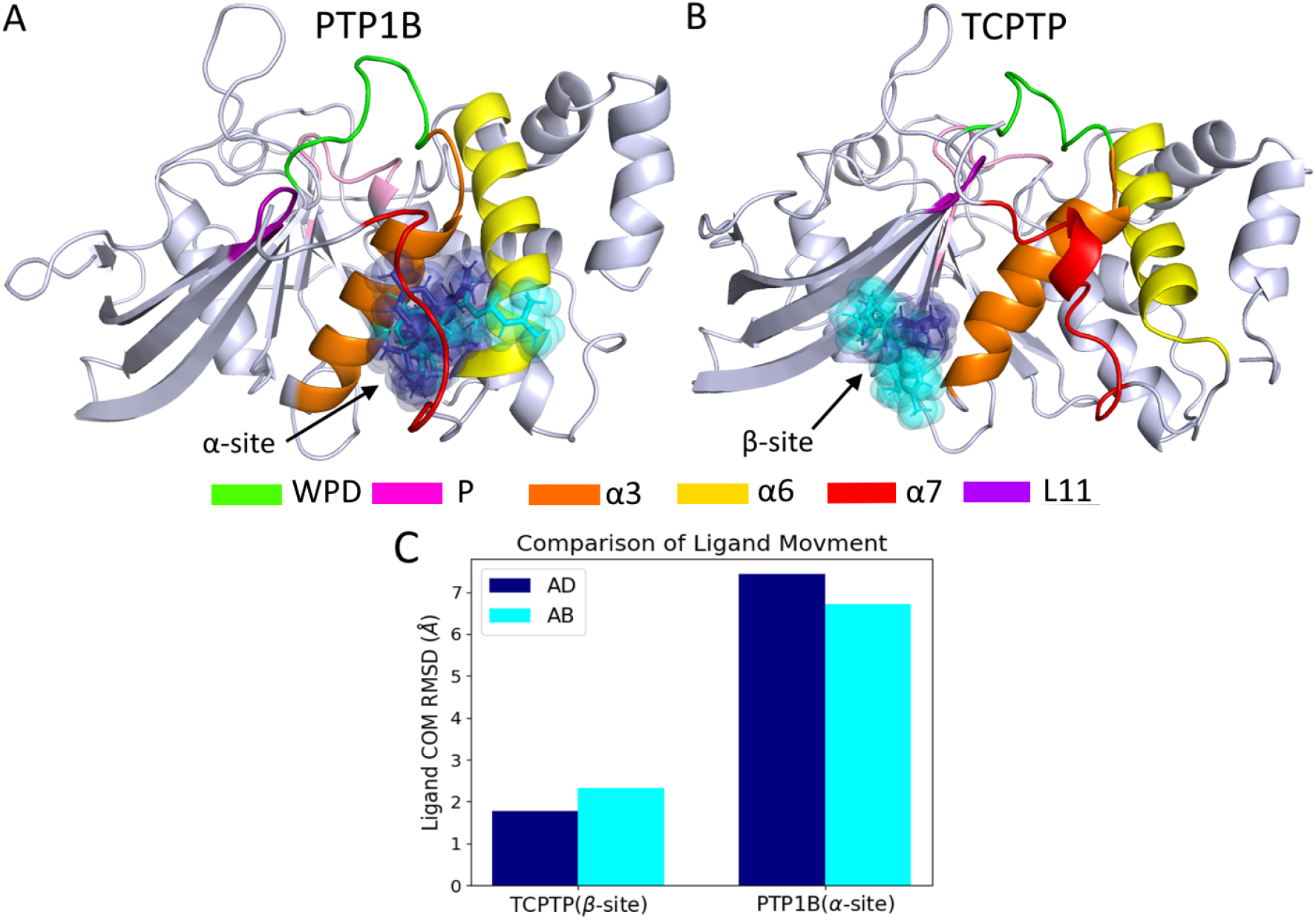
AD and AB bind to different sites on PTP1B and TCPTP. (A) When AD (blue) and AB (cyan) bind to PTP1B with a disordered *α*7 helix they primarily bind to the *α* site, where they interact with the *α*3 and *α*7 helices. (B) When the same two ligands bind to TCPTP, they bind to the *β* site, where they interact with *β*8–10 and the *α*3 helix. (C) Binding to the *α* site results in significantly higher COM RMSD than binding to the *β* site because the ligand samples two regions of this site. One is located closer to the *α*6 helix; the other, closer to the *α*3 helix. The *β* site has a single dominant, though somewhat diffuse, binding mode.

In TCPTP, where allostery is less well studied, the h-bond network revealed by our MD simulations is consistent with published biochemical data on the E295A mutant, which has a reduced catalytic activity (*24*). In brief, E295 participates in a h-bond between the *α*3 and *α*7 helices, where disruption should decrease activity. The high similarity between the allosteric networks of PTP1B and TCPTP is also consistent with the similar decrease in activity observed when the *α*7 helix is removed from these enzymes (4-fold for TCPTP (*24*) and 3-fold for PTP1B (*37*)), in addition to prior work suggesting that allosteric communication is a conserved feature of PTPs (*10*).

The allosteric networks of PTP1B and TCPTP also contain non-bonded interactions that contribute to intramolecular communication. A cluster of van der Waals, h-bond, and salt bridge interactions near the C-terminus, for example, stabilizes the ordered *α*7 helix and allows the h-bond network to form. When we compare these interactions between open and closed protein conformations, PTP1B shows an increased concentration of disrupted interactions at the *α*3–*α*7 interface, while TCPTP shows an increased concentration at the L11–*α*7 interface (Figure 1B and S8). Though the disrupted interactions are different between the two allosteric networks, they connect the same functional protein regions and, thus, appear to exhibit a similar allosteric functionality between PTP1B and TCPTP. Notably, for PTP1B, both interaction interfaces are disrupted by binding to the allosteric site (*3*), an indication that they are unlikely to contribute to differences in inhibitor potency at that site.

### Allosteric ligands bind to a different site in TCPTP than PTP1B

The similarity of the allosteric networks in PTP1B and TCPTP suggests that differences in the potency of nonpolar terpenoid inhibitors for these two enzymes results not from differences in allosteric communication with the active site, but rather from differences in the binding process itself. Our analysis of binding locations supports this interpretation. When bound to PTP1B, AD and AB bind to an *α* site located between the *α*3 and *α*7 helices (Figure 2A). By contrast, when bound to TCPTP, both ligands bind a separate, distinct *β* site that sits at the interface of sheets *β*8–*β*10 and the *α*3 helix (Figure 2B). When these ligands are initialized at the *α* site of TCPTP they move to the *β* site within 100 ns, an indication that the *α* site does not permit the formation of a stable complex with TCPTP. Intriguingly, the two sites permit different amounts of ligand mobility: when bound to the *α* site on PTP1B, AD and AB sample two neighboring sites and, thus, have a larger COM RMSD than when bound to the *β* site of TCPTP (Figure 2C). When bound to the *β* site of TCPTP, AB is more flexible than AD, but both ligands occupy the same binding location and maintain conserved non-bonded interactions with TCPTP (Figure S2).

Binding to the *β* site on PTP1B is still possible, but with reduced binding affinity. In our previous analysis, we simulated the PTP1B-AD complex with a truncated *α*7 helix; for this truncation variant, AD occupied the *β* site (termed loc4 in this previous study) for 88% of observed trajectories compared to 0% occupancy in trajectories with a disordered *α*7 helix (*3*). This truncation-dependent binding is consistent with weaker binding to the *β* site, where residues are conserved between PTP1B and TCPTP. In kinetic assays, the potency of AD for the *α*7-less form of PTP1B was similar to its potency for TCPTP (*4*), a reduction in potency that further corroborates our interpretation that the *α*7 helix stabilizes binding to the *α* site on PTP1B and that the secondary binding location is similar to that sampled in the TCPTP-AD complex.

We probed binding to the *β* site further by using a focused mutational analysis. To begin, we modeled these PTP variants with mutations likely to disrupt binding to the *β* site; in selecting these mutations, we avoided charge-altering residues that might disrupt protein folding. In MD simulations, mutations L158F, S147T, S147V, V156S, and V156T reduced ligand retention time at the *β* site of TCPTP, while analogous mutations in PTP1B had no effect on either ligand retention or COM RMSD at the *α* site (Figure 3A and S3). The localized influence of these mutations suggests that their disruptive effect might serve as a diagnostic for detecting binding to the *β* site. We followed up with in vitro kinetic experiments. For both PTP1B and TCPTP, we measured the IC50 of AD, a readily synthesizable ligand with a crystallographically resolvable binding site. First, we examined wild-type and L158F mutants (Figure S4 and S5). As this mutation reduced ligand retention time for the *β* site of TCPTP, we expected that it would reduce binding and weaken inhibition for TCPTP, but not PTP1B. In contrast to our hypotheses, the IC50s for AD were indistinguishable between wild-type and mutant enzymes (Figure 3B). Measurements of inhibition provide only an indirect— and substrate-sensitive— means of detecting changes in inhibitor binding, so there are a range of reasons why we might not see the expected effect, other than simply inaccurate molecular models. The insensitivity could also result from alternate binding conformations or sites that were not sampled in our simulations. The insensitivity could also result from flexibility in the protein backbone which may allow room for both the bulkier PHE residue and the ligand at with rearrangements at a longer timescale than 300 ns, which is plausible given the location of residue 158 on the surface of the protein where it is more capable of adjustments.

**Figure 3:**
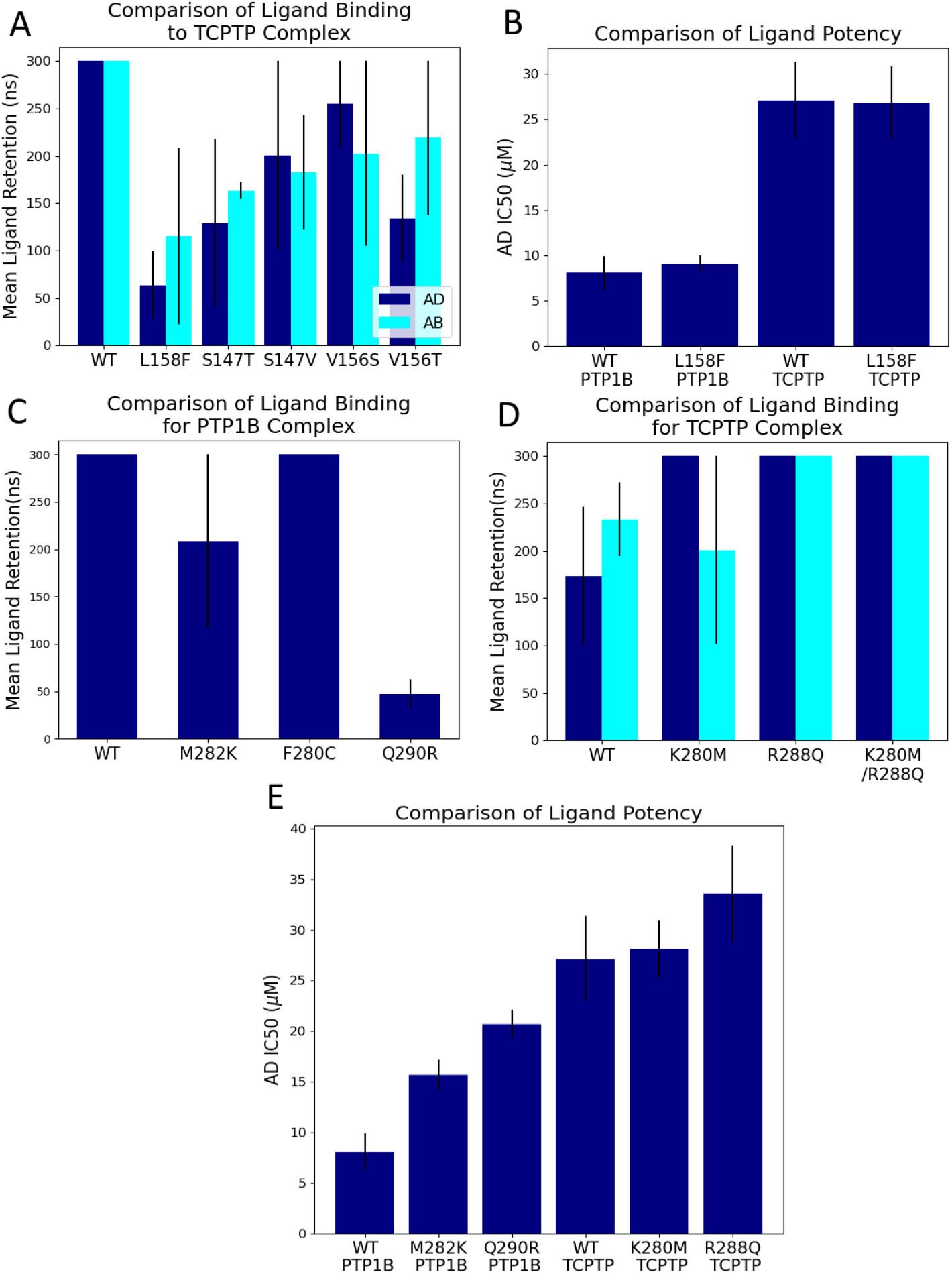
Mutations in the *α* and *β* sites have different effects on the binding of AD to PTP1B and TCPTP.(A) In MD simulations of TCPTP, mutations in the *β* site designed to disrupt binding reduced the mean ligand retention time for both AD and AB. Differences in retention time between mutants were not statistically significant (p < 0.05), a potential limitation of the 300 ns simulations. (B) Experimentally determined IC50 values for AD-mediated inhibition were the same for both wild-type and L158F variants of PTP1B and TCPTP. The insensitivity for PTP1B, where AD binds to the *α* site, is consistent with simulations, but the insensitivity for TCPTP, where L158F reduces ligand retention time in simulations is in contrast to the hypothesis. (C) Mutations that made the *α* site of PTP1B more like the *α* site of TCPTP (M282K, F280C, and Q290R) reduced ligand retention time of AD. (D) Mutations that made the *α* site of TCPTP more like the *α* site of PTP1B (K280M and R288Q), in turn, increased the ligand retention time. (E). Experimentally determined IC50 values for AD-mediated inhibition of PTP1B suggest that M282K and Q290R disrupt inhibition by this compound, a result consistent with simulations, but show that K280M and R288Q in TCPTP have no perceptible effect.

A comparison of residues that line the *α* site of PTP1B and TCPTP indicates that minor sequence differences between PTP1B and TCPTP could destabilize the binding of AD and AB to TCPTP. In particular, prior biophysical analyses suggest that binding to the *α* site of PTP1B can be stabilized by *π*-stacking with PHE280 and PHE196, and through additional non-bonded interactions with the *α*3 and *α*6 helices, primarily residues ALA189, ASN193, and SER295 (Figure S12) (*22*). Of these residues, the *α* site of TCPTP lacks only PHE280, but it also includes several charged amino acids in place of less polar ones found in PTP1B (Figure S9A). These differences should disrupt nonpolar interactions with the *α* site. In discussing this effect, we note that the Amber *ff99sb-ildn* force field used for these MD simulations does not directly incorporate *π*-stacking interactions; however, the Lenard-Jones potential can recapitulate some of the attractive nature of these interactions, with issues primarily arising in the directionality of the interactions which is of less importance for this study.

We performed a similar mutational analysis of the *α* site to understand these effect. Starting with simulations of PTP1B, we introduced analogous residues from TCPTP in the *α* binding pocket. As discussed above, we speculated that the F280C mutation might prevent *π*-stacking with AD and AB, while M282K and Q290R would disrupt nonbonded interactions within the hydrophobic pocket. Only the latter two mutations were sufficiently disruptive to cause ligand dissociation within 300 ns (Figure 3C). The reverse mutations in TCPTP (K280M and R288Q), in turn, increased ligand retention time, an effect consistent with stronger binding at the *α* site (Figure 3D). Although F280C was less impactful in PTP1B, the lack of stabilizing non-bonded interactions did cause AD to move to the top of the pocket (Figure S11), potentially weakening binding. These simulations suggest that F280 helps anchor AD to the bottom of the *α* site but also suggest that disruption of the hydrophobic pocket, rather than the absence of *π*-stacking interactions, is the primary destabilizing factor for ligand binding to this site in TCPTP.

We followed up on our simulations by using kinetic assays to measure the influence of mutations in the *α* site on inhibition by AD. As expected, the M282K and Q290R mutations in PTP1B reduced potency (i.e., increased IC50), an effect consistent with a reduction in binding stability (Figure 3E). The complementary mutations in TCPTP, however, had no effect on IC50 (Figure 3E); that is, they did not yield the expected reduction in IC50 that we might expect from stronger binding to the *α* site. The insensitivity of TCPTP to these mutations suggests that the exchange of a single charged residue in TCPTP to its PTP1B-equivalent may be insufficient to restore full affinity at the *α* site, as the mutations are only suggestive of some level increased binding. Differences in other residues, or in the conformation of the *α*7 helix itself, may also be essential to achieve a detectable increase in binding affinity.

### *β* Site binding destabilizes *α*7

We also found that binding to the *β* site in TCPTP produces distinct disruptive effects in the allosteric network. In PTP1B, binding to the *α* site by both AD and AB disrupts interactions at the *α*3–*α*7 and the L–11–*α*7 interfaces, both of which disorder the *α*7 helix and prevent the formation of a h-bond network that helps stabilize the closed conformation of the WPD loop (Figure4A) (*3*). By contrast, binding to the *β* site of TCPTP disrupts only the L–11–*α*7 interface; in fact, the frequency of several interactions in the *α*3–*α*7 interface increases moderately (<5%) in ligand bound conformations (Figure 4A–B). Accordingly, reordering of the *α*7 helix may only require disruption of the L–11–*α*7 interface; nonetheless, given that the h-bonding network relies on stable h-bonds between the L–11 loop, *α*3, and *α*7 helix disruption of both the *α*3–*α*7 and the L–11–*α*7 interfaces, as seen with binding to the *α* site, may be a more “effective” allosteric disruption.

**Figure 4:**
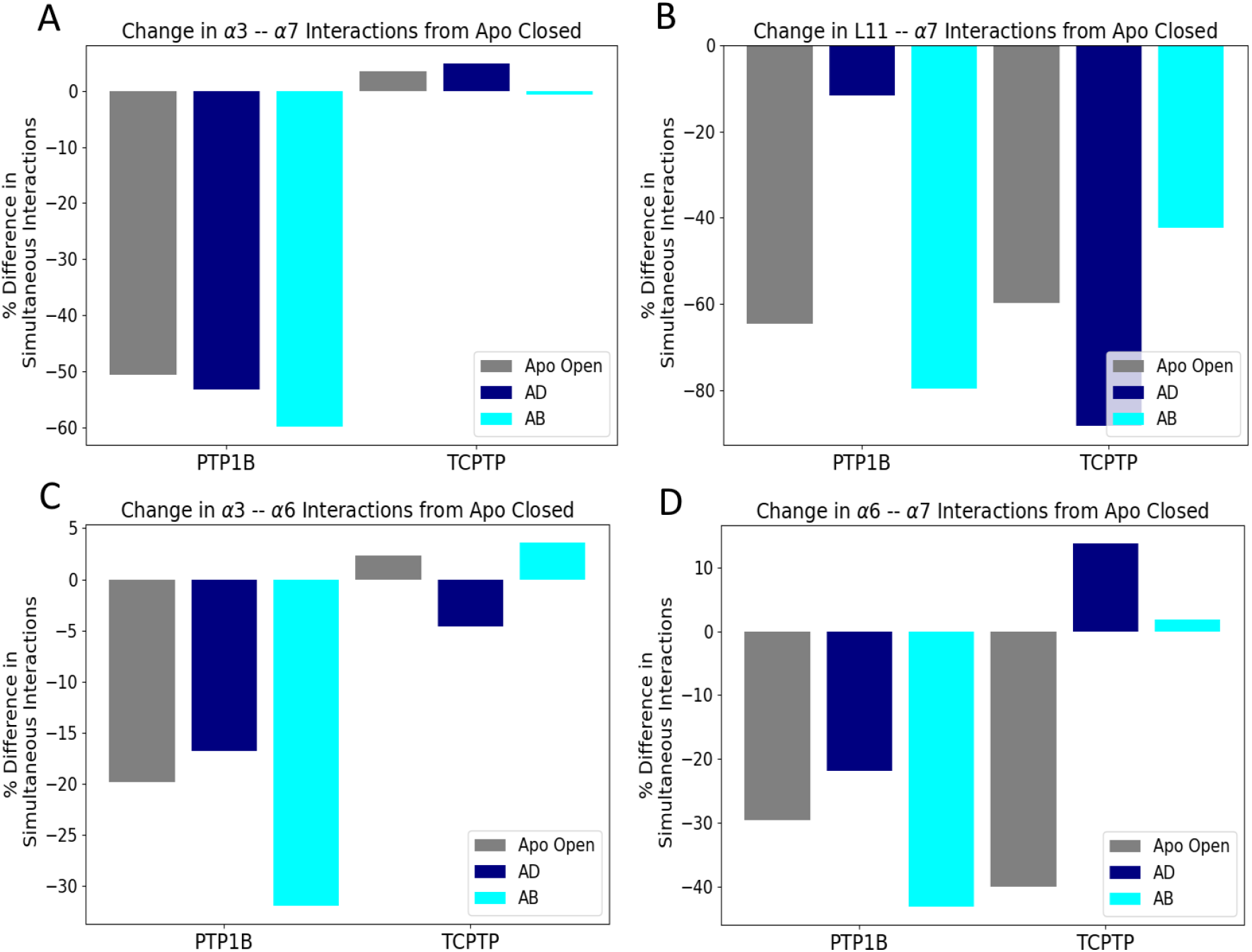
Binding to the *α* and *β* sites yields different disruptive effects. (A) The binding of AD and AB to the *α* site of PTP1B disrupts the *α*3–*α*7 interface. In TCPTP, neither the apo open nor ligand-bound states reduce significantly affect this interface. This plot shows the difference in simultaneous interactions between the apo closed state and each other state. (B) In TCPTP, the apo open, AD-bound, and AB-bound states disrupt the L–11–*α*7 interface; for PTP1B, only the apo open and AB-bound states have a similar disruptive effect. (C-D) For PTP1B, the apo open, AD-bound, and AB-bound states disrupt (C) the *α*3–*α*6 and (D) the *α*6–*α*7 interfaces, where ligand binding to TCPTP has little effect. In previous studies, these interfaces did not appear to have significant effect on the h-bond network, thus the differences between PTP1B and TCPTP are likely to be less important.

## Discussion

Achieving selective inhibition within a highly conserved class of enzymes is difficult and requires extensive structural knowledge of both onand off-targets, and their known inhibitors. The homologous enzymes PTP1B and TCPTP show the same relationship between the WPD loop and the *α*7 helix; when the WPD loop reorients from open to closed, the *α*7 helix folds to ordered conformation from a disordered one. Our simulations support the existence of a conserved allosteric network between the WPD loop and the *α*7 helix and indicate that subtle differences in network residues or peripheral non-bonded interactions are unlikely to contribute to differences in the potency of allosteric inhibitors for PTP1B and TCPTP. Instead, minor residue differences in the *α*6 and *α*7 helices appear to direct inhibitor binding to different sites.

For TCPTP, the hydrophobic pocket that allows for stable binding to the *α* site in PTP1B is disrupted by the distinct charged residues at position 290, and to a lesser extent, position 282. Instead, these ligands bind to a *β* site, which is conserved in both PTP1B and TCPTP, and where prior simulations of PTP1B suggest that binding is likely to occur in the absence of the *α*7 helix. In either protein, binding to the *β* site appears to be less effective at modulating the allosteric network. Intriguingly, a recent study demonstrates that binding to the *β* site of PTP1B with a truncated *α*7 helix is not limited to fully nonpolar inhibitors (*38*). Our observations suggest that these inhibitors might also bind to the *β* site of TCPTP.

Broadly, our findings highlight the role of the *α*7 helix in enabling selective interactions, and provided a framework for tuning the selectivity of inhibitors through focused interactions at the *α* and *β* sites. Many promising allosteric inhibitors of PTP1B have large nonpolar surface areas and could exhibit similar binding behavior to TCPTP (*39* – *41*). The conserved allosteric network of PTP1B and TCPTP indicates that allosteric modulation of both proteins is possible; however, the results of the present study suggest that binding to the *α* site is likely to be more selective for PTP1B. The design of selective inhibitors for TCPTP, in turn, may be more challenging, given the similarity of the *β* sites; however, compounds that exploit charged residues within the *α* site, or that interact with unique residues in the disordered region following the *α*7 helix, where sequence conservation is low, could offer a path forward (*42*).

## Author Contributions

A.J.F, J.M.F. and M.R.S conceptualized the project. A.J.F. and M.R.S. designed the computational methodology, and E.T.L, L.K., G.W.D., and J.M.F. designed the experimental methodology. A.J.F. and H.M.P. performed and analyzed all molecular simulations experiments. A.J.F. wrote the original draft. H.M.P., J.M.F. and M.R.S. edited and reviewed the manuscript. L.K., G.W.D., and E.T.L. expressed and purified the proteins. L.K. performed molecular cloning and in vitro kinetic assays. J.M.F. and M.R.S. supervised the project and obtained the resources.

## Acknowledgements

This work was supported by funds provided by the University of Colorado Boulder (A.J.F, M.S., and H.M.P), the National Institute of General Medical Sciences of the National Institutes of Health (E.T.L., R35GM143089), the National Science Foundation (J.M.F., CBET 1750244), and Think Bioscience (L.K.). This work utilized computational resources from the University of Colorado Boulder Research Computing Group, which is supported by the National Science Foundation (awards ACI-1532235 and ACI-1532236), the University of Colorado Boulder, and Colorado State University. This work also used the Advanced Cyberinfrastructure Coordination Ecosystem: Services & Support (ACCESS), which is supported by National Science Foundation grant number ACI-1548562. Specifically, it used the Bridges-2 system, which is supported by NSF ACI-1928147 at the Pittsburgh Supercomputing Center (PSC).

