## Supplementary Information for "A biophysical rationale for the selective inhibition of PTP1B over TCPTP by nonpolar terpenoids"

| Component | Organism | Plasmid | Source |
|---|---|---|---|
| PTP1B | *H. Sapiens* | pET21B_PTP1B | Nicholas Tonks, Cold Spring Harbor |
| TCPTP | *H. Sapiens* | pGB100_TCPTP | Addgene: 33365 |

Table S1: Gene Sources

| Mutant | F Primer | R Primer |
|---|---|---|
| PTP1B (L158F) | CAAGTCATATTATACAGTGCG ACAGttcGAATTGGAAAACC TTACAACCCAAGAAAC | CTGTCGCACTGTAT AATATGACTTGATATC |
| PTP1B (M282K) | GTGATCGAAGGTGCCAAATTCA TCaaaGGGGACTCTTCCGTGCAG | GATGAATTTGGC ACCTTCGATCAC |
| PTP1B (Q290R) | GGGACTCTTCCGTGCAGGATcgt TGGAAGGAGCTTTCCCACGAGG | CCTGCACGGAA GAGTCCCCC |
| TCPTP (L158F) | GATGTGAAGTCGTATTATACA GTACATttcCTACAATTAGAA AATATCAATAGTGGTG | GTACTGTATAATACG ACTTCACATCTTCTGAC |
| TCPTP (K280M) | CTATAATAGAAGGAGCAA AATGTATAatgGGAGATT CTAGTATACAGAAACGATG | CATTTTGCTCCTTC TATTATAGCCATGTATG |
| TCPTP (R288Q) | GTATAAAGGGAGATTCTAG TATACAGAAAcagTGGAAA GAACTTTCTAAGGAAGAC | CTGTATACTAGAATCT CCCTTTATACATTTTGCTC |

Table S2: Primers used for introducing point mutations

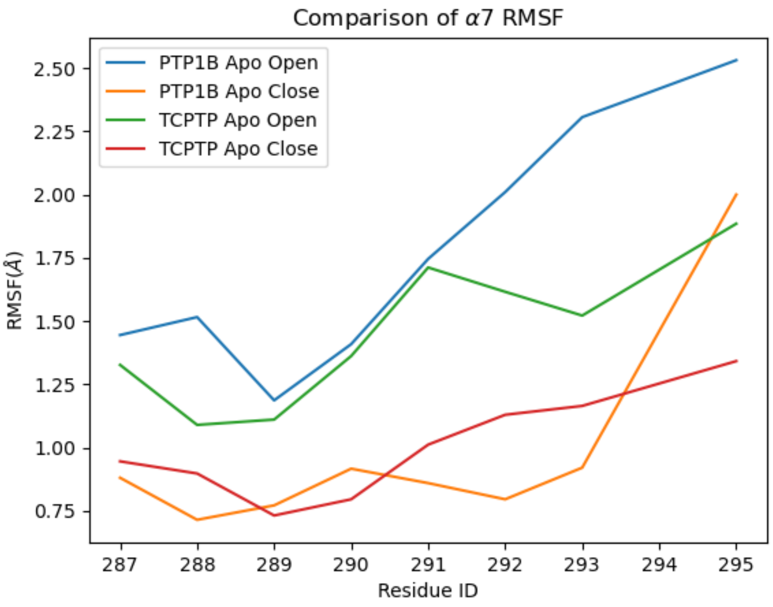


Figure S1: Despite the fact that the $\alpha7$ helix on TCPTP has fewer stable non-bonded interactions, the overall flexibility as shown by the RMSF of residues in the $\alpha7$ helix is not significantly different from that of corresponding the apo PTP1B trajectories.

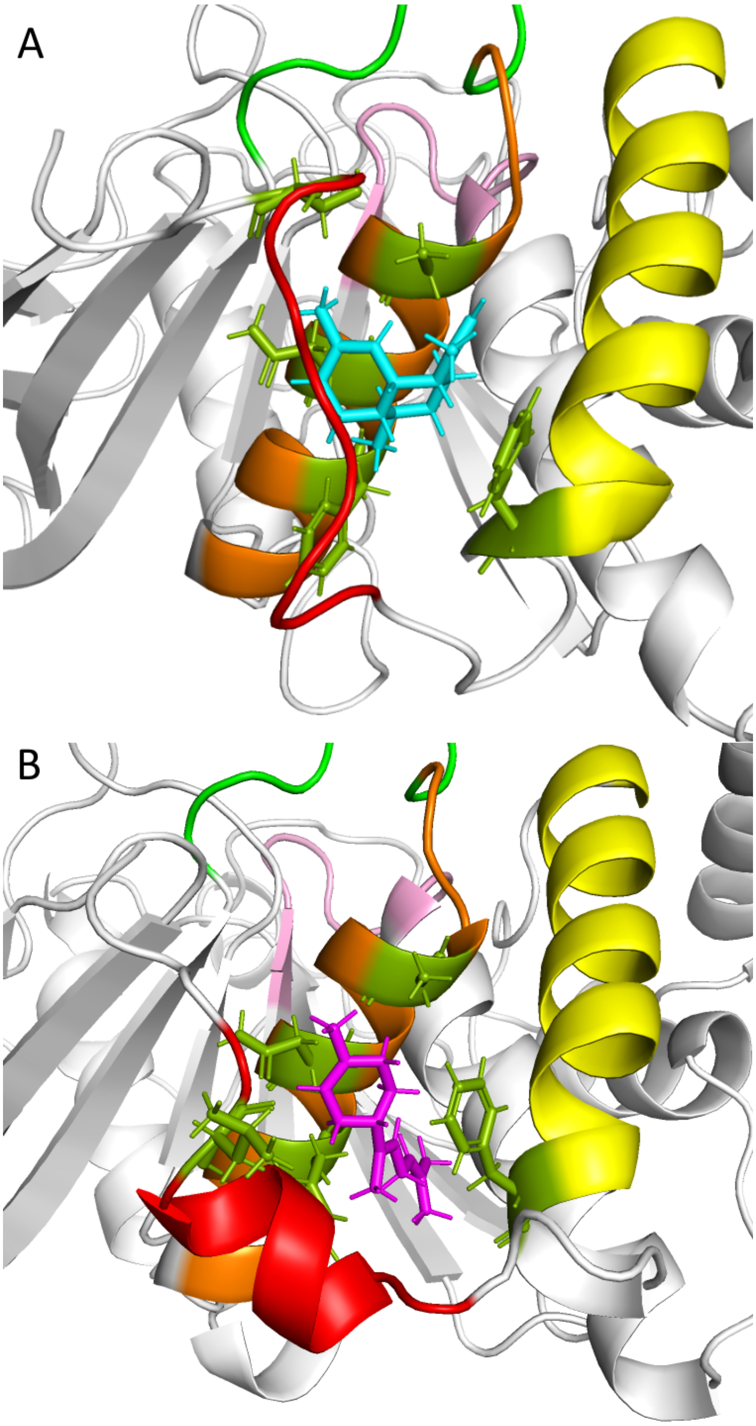


Figure S2: The centroid clustered on ligand heavy atoms for trajectories of PTP1B in complex with AD (A) and AB (B). Both ligands form nearly identical non-bonded interactions with residues shown in green. The only difference in residue binding is that AD forms more frequent non-bonded interactions with residue 295 in the $\alpha$7 helix while AB forms more frequent interactions with residue 294. These differences are likely due to flexibility in the $\alpha$7 helix leading to small differences in which stable non-bonded interactions form with the ligand.

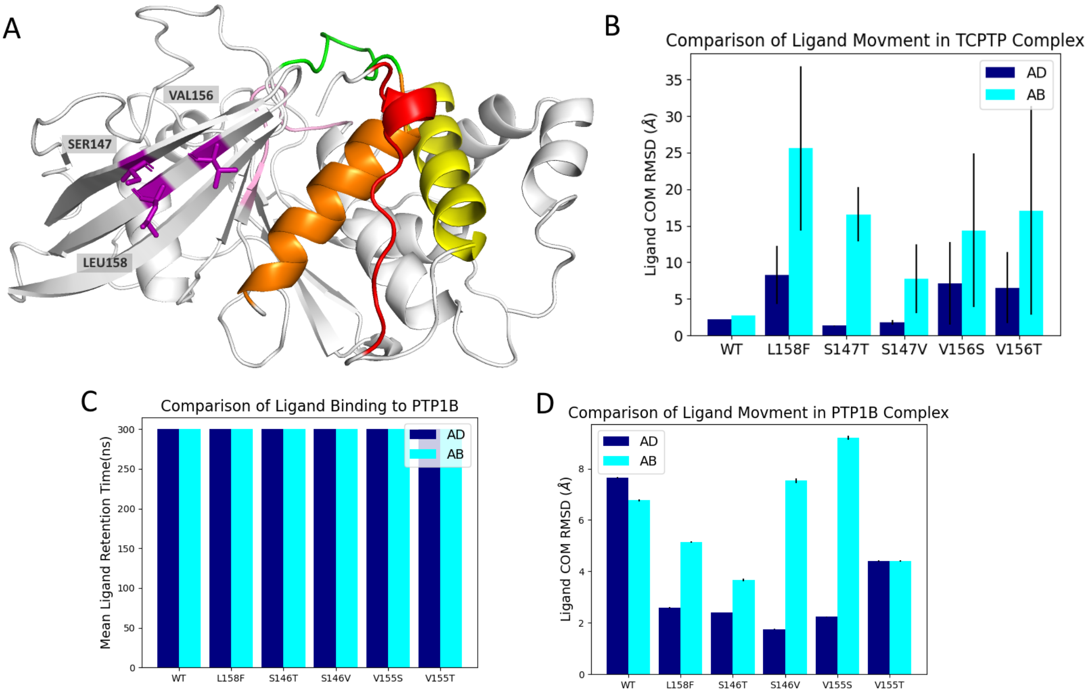


Figure S3: (A) The three residues which were mutated are the primary points of contact with the ligand when bound to the $\beta$-site on TCPTP. (B) Because of this limitation we also evaluated the relative ligand flexibility when bound to the $\beta$ site for each mutant using the COM RMSD. The increase in the COM RMSD of the ligands even when remaining bound to the mutated $\beta$ site shows the destabilization of the binding site before the ligand dissociates completely. Mutants which disrupt binding to the $\beta$ site do not disrupt binding to the $\alpha$ site as shown by the ligand retention time (C) and COM RMSD values (D) for the ligands bound at the $\alpha$ site of PTP1B for WT compared to mutants.

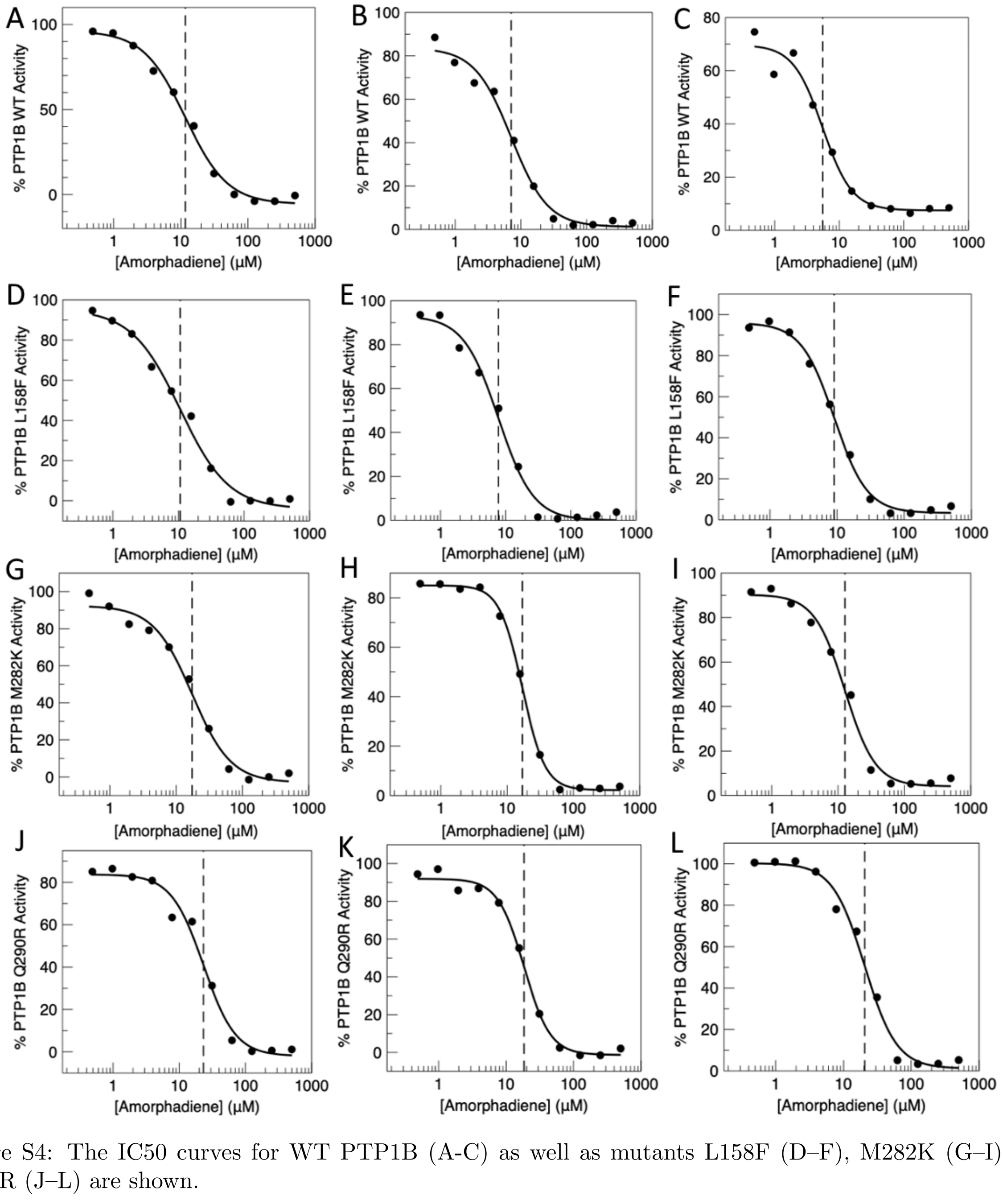


Figure S4: The IC50 curves for WT PTP1B (A-C) as well as mutants L158F (D–F), M282K (G–I), and Q290R (J–L) are shown.

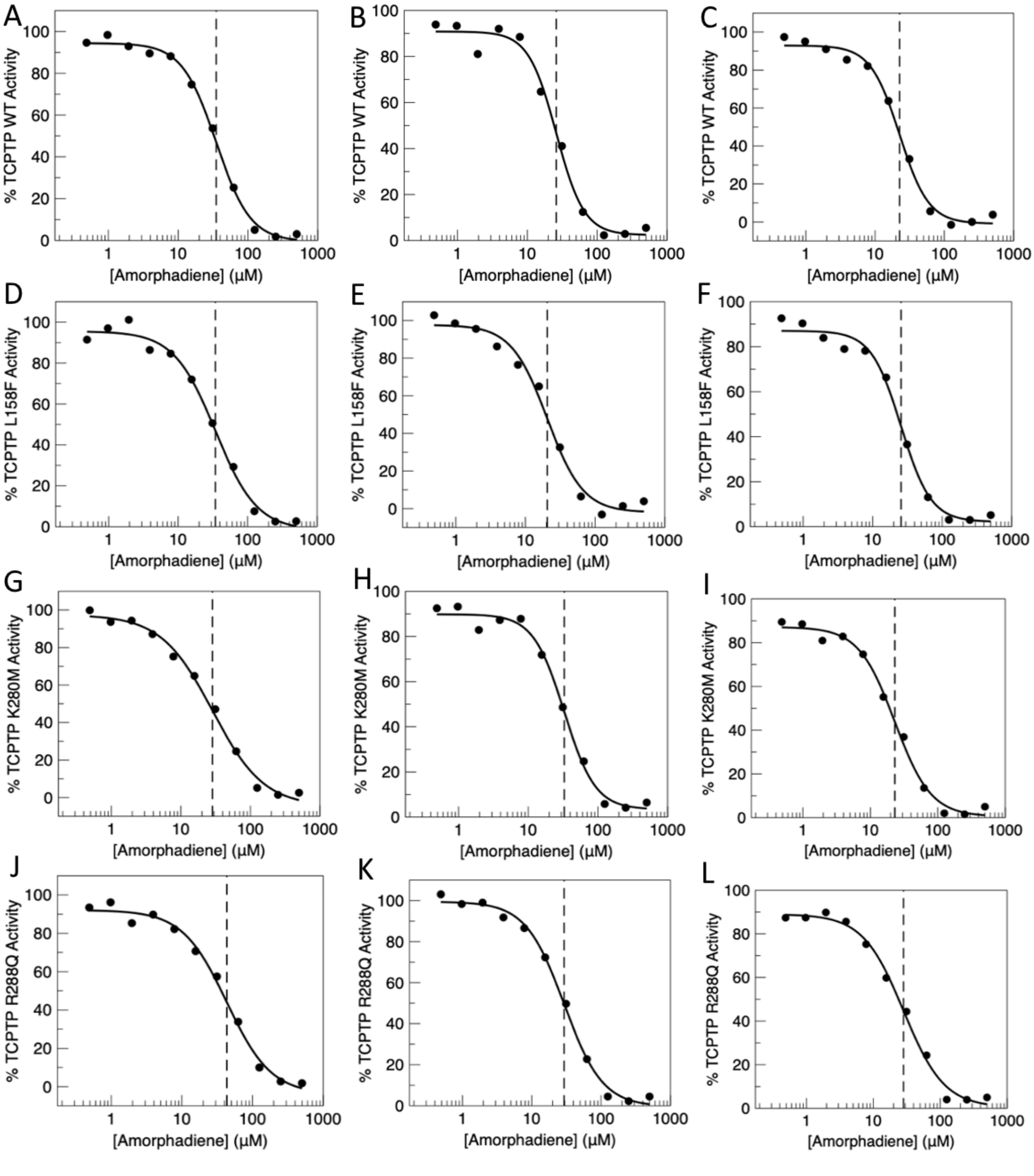


Figure S5: The IC50 curves for WT TCPTP (A-C) as well as mutants L158F (D–F), K280M (G–I), and R288Q (J–L) are shown.

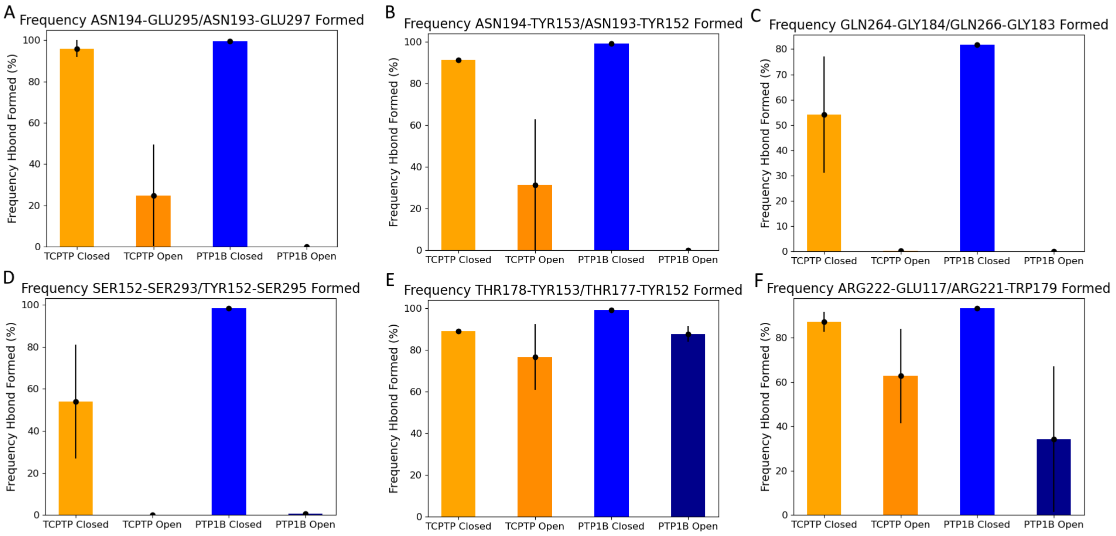


Figure S6: Core bonds central to the h-bond network in both PTP1B and TCPTP. Each bar shows the mean bond occupancy percentage from 3-4 trajectories in either the apo open or closed state for each protein. The error bars show the standard error on the mean for each set of trajectories. Given the difference in residue numbering between TCPTP and PTP1B the bond names are shown as "TCPTP bond name/PTP1B bond name." A–B have high error bars for the TCPTP open state due to a single outlier trajectory which showed high occupancy in both of these bonds. This outlier trajectory also had the highest mean $\alpha$ helicity for the $\alpha$7 helix suggesting which is likely a significant contributing factor. In D, the bonds compared differ significantly in residue numbering as in PTP1B the bond connects the P-loop to the WPD loop while in TCPTP the P-loop is connected to the substrate binding loop which is adjacent to the WPD-loop in the active site. These bonds were compared directly as they both position the catalytic cystine in a similar manner despite connecting distinct structural elements.

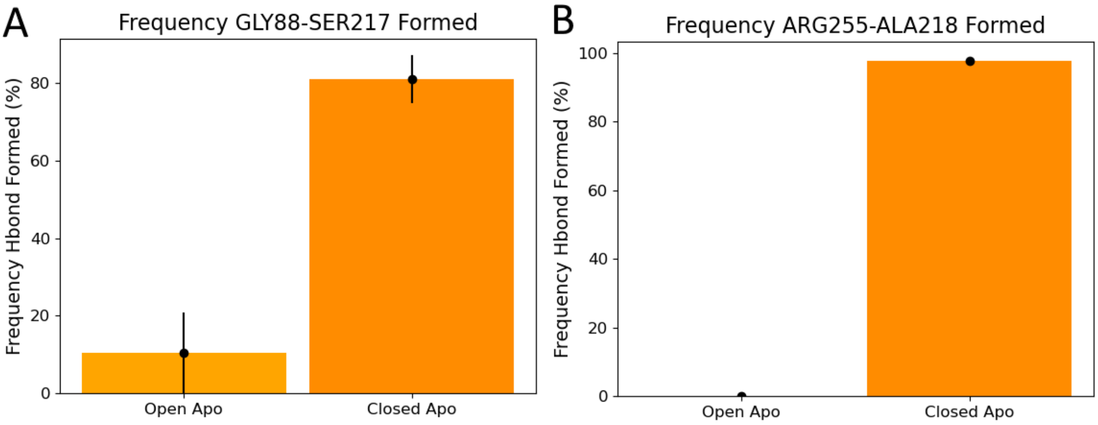


Figure S7: These additional bonds were present in the TCPTP allosteric network but not in that of PTP1B. Both bonds are formed between the P-loop and other protein regions which allow for proper positioning of the catalytic cysteine within the active site. The presence of such bonds does not appear to affect the allosteric network as the P-loop does not undergo significant conformational changes between the apo open and closed conformations.

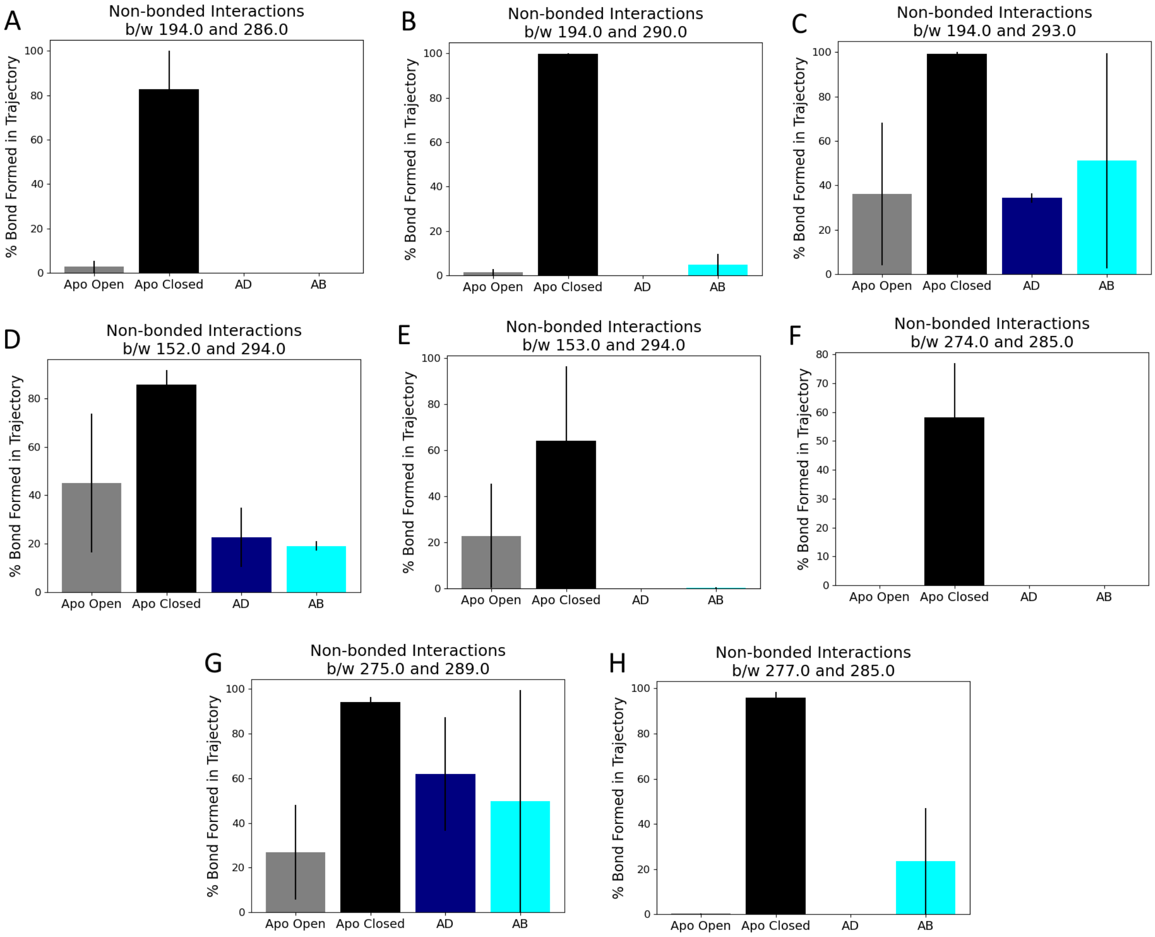


Figure S8: As TCPTP transitions from the apo open to apo closed conformations a series of non-bonded interactions break between the $\alpha 7$ helix and the $\alpha 3$ helix and L–11 loop. This figure shows the individual interactions which are disrupted during this conformational change at the $\alpha 3$–$\alpha 7$ (A–C), L11–$\alpha 7$ (D–F), and $\alpha 6$–$\alpha 7$ (G–H) interfaces.

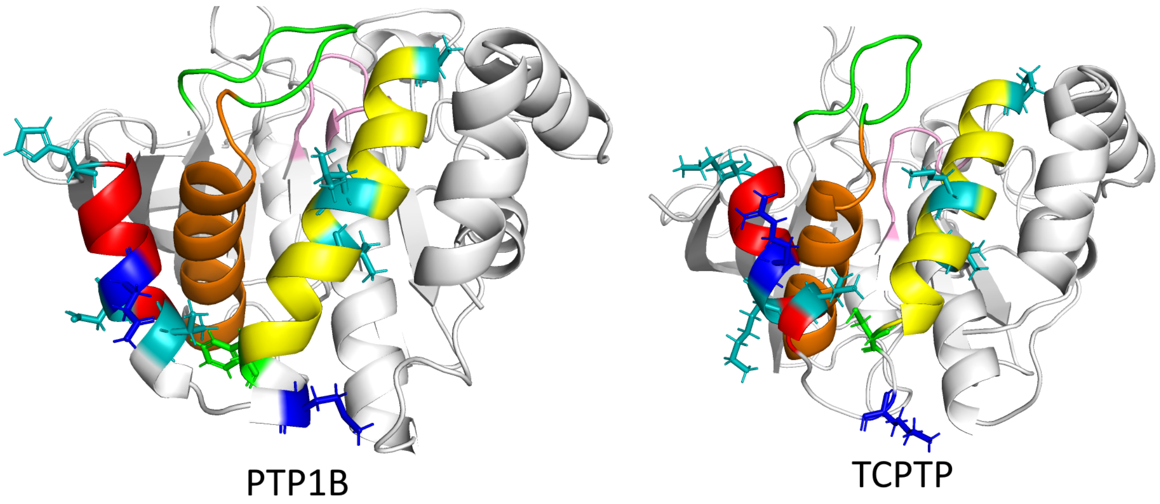


Figure S9: PTP1B and TCPTP have high sequence similarity and nearly identical secondary and tertiary structure. Above a sequence comparison within the allosteric site shows that of the nine residues which are distinct between PTP1B and TCPTP in both protein structures. In teal are residues which have moderate changes in size or change charge. In blue are residues which are non-polar in PTP1B but are charged in TCPTP and in green is residue 280 (278 in TCPTP) which loses the ability to form $\pi$-stacking interactions.

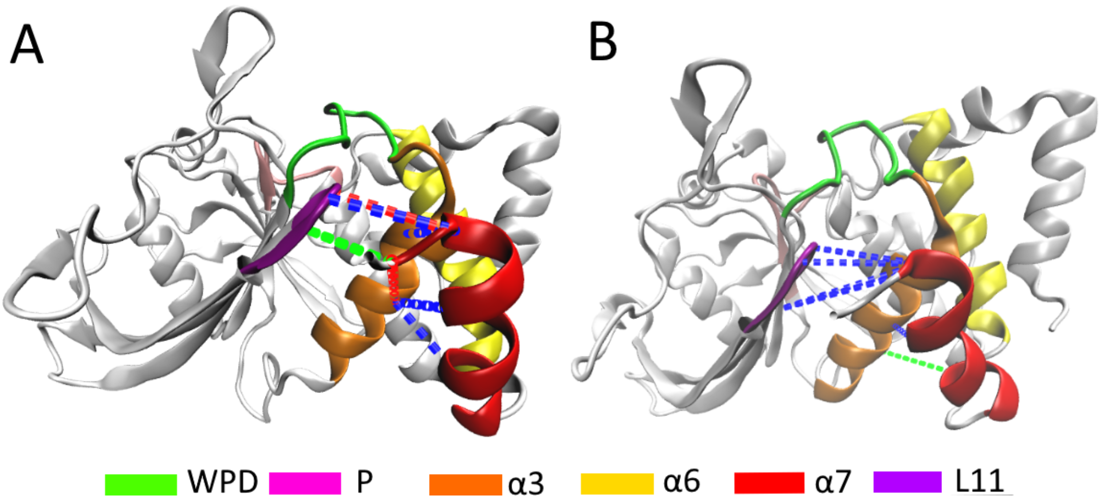


Figure S10: Non-bonded interactions which are consistently disrupted in the open state for PTP1B (A) and TCPTP (B). The non-bonded interactions between helices fluctuate so in any frame there are more interactions which are disrupted than shown here but this is limited to those interactions which are individually present significantly (p<0.05) less in the open vs closed state. No interactions at the $\alpha3 - \alpha6$ interface fit this criteria and thus none are shown. H-bonds are labeled in black, ionic interactions are labeled in green, and all other non-bonded interactions in blue. Overall these interactions demonstrate how the disordering of the $\alpha7$ helix is primarily enabled by disrupting the L11–$\alpha7$ interface for TCPTP while in PTP1B these disrupted interactions are more evenly distributed between the L11–$\alpha7$ and the $\alpha3$–$\alpha7$ interfaces. Allosteric inhibitors are thought to primarily prevent the reordering of the $\alpha7$ helix and thus the disrupted interface has potentially significant implications for the allosteric modulation of the inhibitors. However, the conservation of the h-bond network suggests any difference in allosteric modulation would be minimal.

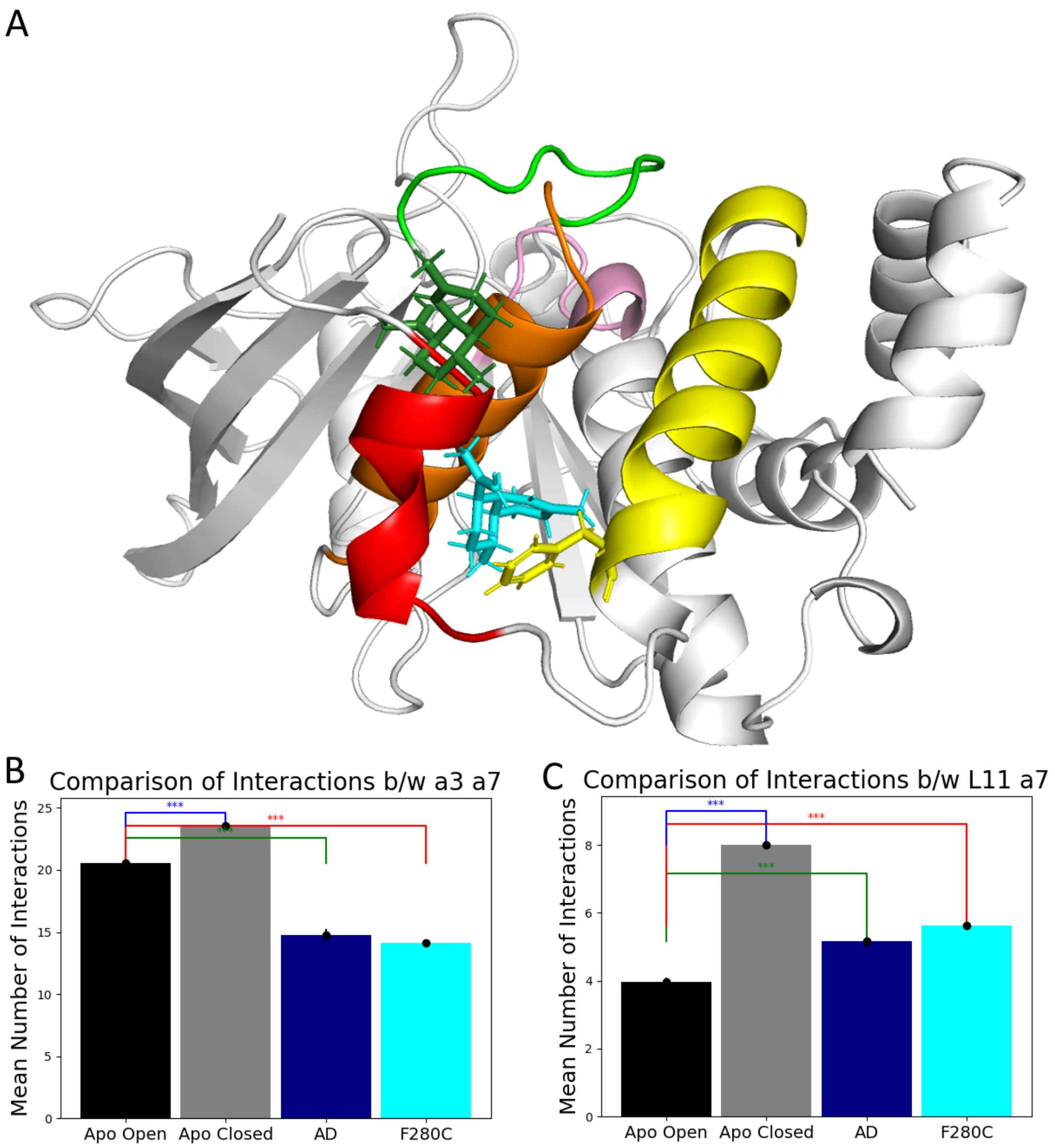


Figure S11: Though the F280C mutation did not cause AD to dissociate from the allosteric site, ligand binding position and orientation was significantly changed leading to differences in the potency of the inhibitor. (A) In WT PTP1B (blue), AD binds near the connection between the $\alpha 6$ and $\alpha 7$ helices where it is anchored by F280 residue interactions. In the F280C mutant (green), AD moves to an alternate position which forms no non-bonded interactions with the $\alpha 6$ helix. Overall the ligand forms few stable non-bonded interactions suggesting the ligand has a considerable decrease in affinity for this binding location. When bound to F280C, AD does still disrupt interactions in a similar manner primarily at the $\alpha 3$–$\alpha 7$ interface and to a lesser extent at the $\alpha 7$–L11 interface (B).

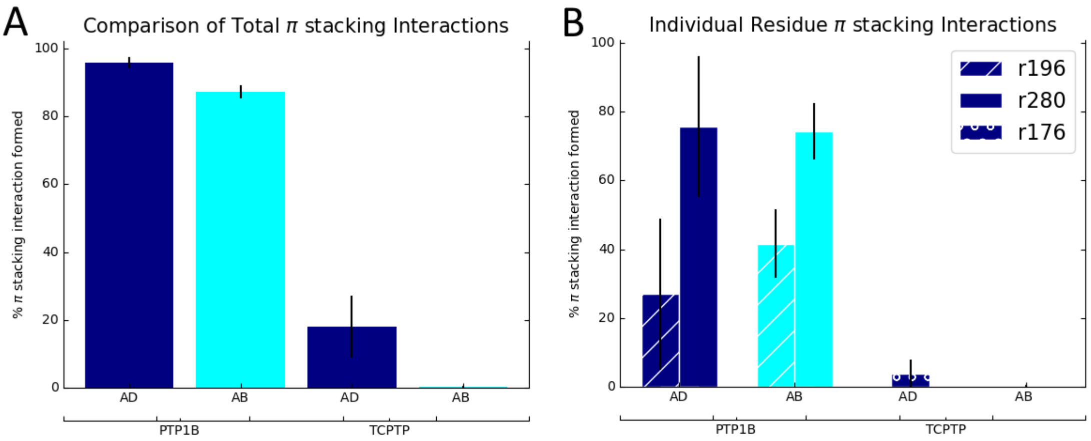


Figure S12: Allosteric inhibitor binding to PTP1B is theorized to be stabilized by the formation of pi-stacking interactions with PHE280. (A) This figure depicts the percent of the trajectory for which any $\pi$-stacking interactions are present between the protein and ligand when in complex with both PTP1B and TCPTP. It is clear that while PTP1B forms stable $\pi$-stacking interactions with both ligands TCPTP does not form these interactions. (B) In PTP1B binding to either AD or AB the $\pi$-stacking interactions with the ligand are primarily with residue PHE280 although there are fleeting $\pi$-stacking interactions with PHE196. While PHE280 is not conserved in TCPTP, PHE196 is, but we do not observe $\pi$-stacking interactions with this residue. There are fleeting $\pi$-stacking interactions with PHE176 in TCPTP, but they are not well conserved or stable.
